# Layer 1 NDNF^+^ Interneurons Control Bilateral Sensory Processing in a Layer-dependent Manner

**DOI:** 10.1101/2021.12.02.470902

**Authors:** Rasmus Vighagen, Lorenzo Gesuita, Angeliki Damilou, Anna Cavaccini, Lila Banterle, Veerle Steenhuis, Theofanis Karayannis

## Abstract

Bilateral sensory information is indispensable for navigating the world. In most mammals, signals sensed by either side of the midline will ultimately reach the cortex where they will be integrated for perception and appropriate action selection. Even though information transferred across the hemispheres is routed through the corpus callosum, how and which microcircuits are key in integrating it is not well understood. Here we identify an essential role for layer 1 NDNF^+^ inhibitory cells of mice in integrating bilateral whisker-evoked information in an NMDA receptor-dependent manner. Direct connections from the contralateral cortex and the ipsilateral side activate NDNF^+^ neurons, which subsequently inhibit the late spiking activity of underlying layer 2/3 neurons, but not layer 5. Our results identify a feed-forward regulatory pathway for bilateral cortical sensory processing of upper layer cortical neurons actuated via layer 1 NDNF^+^ interneurons.

## Introduction

Many animals use a symmetrical pair of sensory receivers they possess on either side of the midline in order to explore aspects of their environment. The bilateral information provided by all sensory modalities enable the perception of a three dimensional world. For this to happen the information coming from both sides of the body has to be integrated, a process that takes place within the cerebral cortical circuits. A set of bilateral sensory organs that mice rely on the most for spatial navigation and object recognition are their whiskers. These facial hairs receive tactile stimuli that are transmitted via ascending pathways that sequentially encompass the brainstem, the thalamus and finally an area within the cortex called the vibrissae primary somatosensory cortex (vS1), or alternatively ‘barrel cortex’. There is a barrel cortical area within each hemisphere, processing inputs detected from the contralateral whisker pad. Nevertheless, activity reaching one hemisphere is also passed over to the opposite one, through callosal projections of pyramidal cells of the supra- and infra-granular layers (Petreanu et al. 2007). Such contralaterally-projecting neurons in vS1 have been shown to target both homotopic-, reciprocally-connected- and non-reciprocally-connected areas, such as the secondary whisker area of the contralateral side (Jones & Powell 1968). These interhemispheric connections have been shown to be key for synchronization between the two hemispheres in animals (Engel et al. 1991) and for fine motor coordination in humans (Goldman & Nauta 1977, Mihrshahi 2006). Research has uncovered that such callosal connections are not only engaging excitatory, but also inhibitory cells (Carr & Sesack 1998, Cissé et al. 2003, Karayannis et al. 2007). The functional significance of these connections has been demonstrated *in vivo* by delivering sensory stimuli to the hind paw of mice which travel through the thalamocortical projections to the contralateral cortex and then cross over to the ipsilateral cortex via the corpus callosum. This information then reaches the distal dendrites of pyramidal cells of cortical layer 5 (L5) (Palmer et al. 2012). The same study also demonstrated that the stimuli engage inhibitory cells leading to a gamma aminobutyric acid beta receptor (GABA*_B_*R)-dependent control of the L5 pyramidal cell dendritic activation and cell output. However, the inhibitory cells responsible for this phenomenon have not yet been revealed. There is a plethora of GABAergic neuron types distributed throughout the cortical laminae, which are more heterogeneous than previously thought (Tasic et al. 2018). Even though it had been anatomically and functionally described that inhibitory cells receive excitatory inputs from the contralateral side (Carr & Sesack 1998, Karayannis et al. 2007), more recent work using Channelrhodopsin-based connectivity mapping has found that different types of unidentified GABAergic interneurons (INs) in L1, L2/3 and L5 receive excitation from the contralateral side, with L1 neurons displaying the strongest sustained activation (Palmer et al. 2012). Recent work has shown that the diversity of L1 GABAergic cells can be parsed out into four different IN types, the Neurogliaform cell (NGFC), Canopy, α7-nAChR positive (+) and a type of Vasoactive-intestinal-peptide expressing cell (VIP^+^) (Schuman et al. 2019). This study convincingly showed that both NGFCs and the Canopy INs are positive for the neuron derived neurotrophic factor (NDNF), a marker exclusively labelling neurons of adult cortical L1 (Abs et al. 2018). Nevertheless, these two cell types have a number of distinguishing features, both in terms of molecular expression as well as function. The molecular marker Neuropeptide Y (NPY) is only expressed in NGFCs, which have a late-spiking (LS) onset, while Canopy cells do not express NPY and are non-late-spiking (non-LS) neurons (Schuman et al. 2019). Both these neuronal types have been shown to generate a very slow GABA*_A_*R-dependent inhibitory responses to connected pyramidal cells, although in contrast to the Canopy Ins, NGFCs additionally activate GABA*_B_*Rs (Schuman et al. 2019). This leaves NGFCs as the sole L1-residing GABAergic cell type currently described to activate GABA*_B_*Rs even after a single action potential (AP) (Price et al. 2008). Despite the described work on L5 pyramidal cells, it is currently unknown how L2/3 pyramidal cells are engaged in contralateral information transfer and importantly whether such information would be regulated by feed-forward inhibition. This would provide important insights into how the upper layers of the cortex participate directly via interhemispheric connections in higher order cortical processing of bilateral sensory stimuli.

In this study we tested the hypothesis that NDNF^+^ INs residing in L1 are imbued with the key characteristics to integrate bilateral signals and efficiently regulate the excitatory drive onto the dendrites, and ultimately impact the output, of L2/3 pyramidal cells. Our hypothesis rests on the fact that these NDNF^+^ cells bear an elaborate axonal ramification in L1, as well as an unusually large NMDA receptor (NMDAR)-dependent component in their glutamatergic synapses. NMDA receptors are well known for their crucial function as synaptic coincidence detectors and for the non-linear integration of incoming neuronal activity (Chittajallu et al. 2017, Cull-Candy & Leszkiewicz 2004, Price et al. 2008).

Using *in vivo* silicon probe recordings in combination with whisker evoked stimulation, we find that removing NMDA receptors from NDNF^+^ L1 neurons leads to a significant increase in the late AP spiking of L2/3 pyramidal cells, and that this effect is stronger when both whiskers are simultaneously deflected. Probing the observed effect in L2/3 neurons, we find that their modulation is mediated through GABA*_B_*Rs, which indeed points at input originating from NDNF^+^ NGFCs. Surprisingly, by performing *in vivo* whole-cell recordings from L2/3 pyramidal neurons we identify that NDNF^+^ INs are the main providers of contralaterally-driven feed-forward inhibition onto L2/3 cells.

## Results

### NDNF^+^ vS1 neurons are confined to L1, extensively develop in the third postnatal week and receive input from the contralateral vS1

To target the outermost lamina and to assess if L1 NDNF^+^ INs participate in the integration of somatosensory signals and modulate the underlying laminae activity, we utilized an NDNFCre line (denoted as CTRL; Ndnf-IRES2-dgCre-D) (Tasic et al. 2016). In line with published research on different parts of the cortex, we find that when performing fate-mapping of the NDNFCre^+^ cells using a tdTomato reporter mouse line (Ai14), the fluorescent neurons are almost exclusively found within L1 of vS1 (Figure 1a) (Abs et al. 2018, Schuman et al. 2019). This observation matches the mRNA *in situ* hybridization (ISH) for NDNF in images published by the Allen Brain Institute (Atlas. 2004b). Nevertheless, the developmental labelling of NDNF expressing cells with our method also revealed that there are other non-neuronal, vascular system-related cells developmentally expressing the NDNF that are labelled in the cortex, as well as in some subcortical structures. In conclusion our fate-mapping strategy confirms that within the somatosensory cortex, NDNF is a specific marker for L1 neurons, all of which are GABAergic INs (Abs et al. 2018, Schuman et al. 2019). It has recently been shown that L1 NDNF^+^ INs in the auditory cortex receive afferents from ipsilateral cortical sensory areas, as well as from the contralateral cortex (Abs et al. 2018). To test if the latter holds true within vS1, we performed monosynaptic rabies tracings onto the NDNF^+^ population within this area. We injected a pseudo-typed rabies virus (RV) into the NDNFCre mouse cortex, either previously iontoporated with a Cre-dependent HTB plasmid at P3 or crossed with the Cre-dependent HTB mouse line (see methods) allowing for expression of the TVA receptor and the glycoprotein ‘G’ in NDNF^+^ cells (NDNFCre-HTB). The presynaptic partners directly synapsing onto these L1 INs were marked with mCherry (from the RV). The RV injections were carried out at P14 and the tissue was harvested at P24. Whole brains were cut, processed and imaged through a slide scanner. Similar to Letzkus et al., these experiments revealed that the NDNF^+^ neurons within vS1 receive inputs from both hemispheres (Figure 1b).

**Figure 1:**
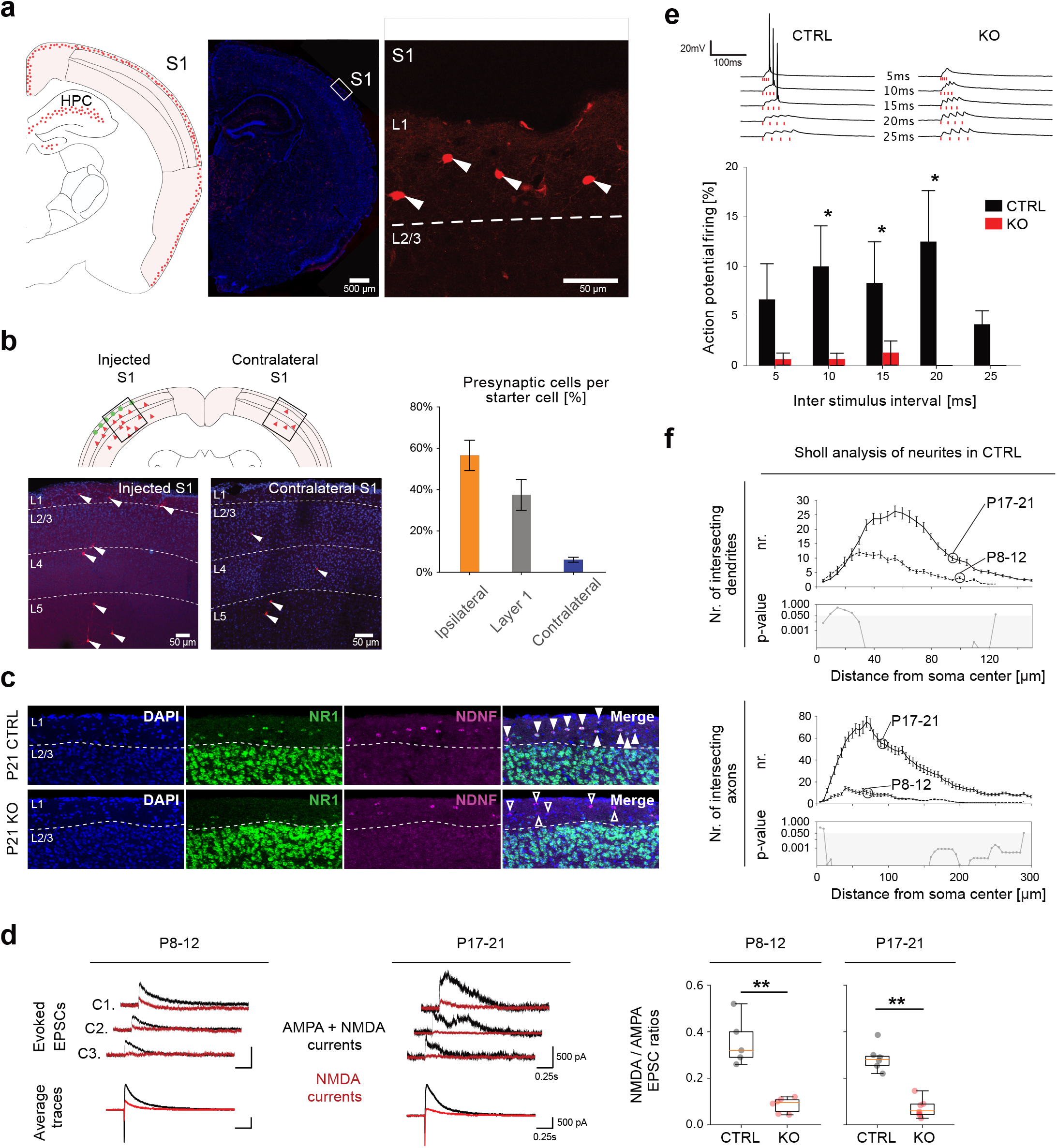
NDNF^+^ vS1 neurons are confined to L1, extensively develop in the third postnatal week and receive input from the contralateral vS1. **(a)** Fate-mapped NDNF^+^ neurons (NDNFCre-Ai14 mice) at P21. Illustration of the position of NDNF^+^ neurons within one hemisphere (left). Coronal image of a whole hemisphere (2^nd^ from left), also delineating the zoom-in region (box) shown on the right. Zoom-in of L1 of vS1 (3^rd^ from left). **(b)** Monosynaptic pseudo-typed rabies tracings carried out at P14-15 in NDNFCre-HTB mice. The animals were injected at P14-15 and the brains were harvested and analyzed at P24-25. Images taken from rabies-injected site of S1 and contralateral S1 at P24. Quantification as percentage of presynaptic cells per given starter cell, normalised per brain (solid arrowheads point at presynaptic cells). **(c)** ISH for NDNF and NR1 mRNA in L1 neurons of S1 in CTRL and KO tissue at P21 (solid arrowheads point at double positive cells, whereas hollow arrowheads point at NDNF positive only cells). **(d)** AMPAR- and NMDAR-dependent (Gabazine added) or NMDAR-dependent (Gabazine+CNQX added) mediated currents were evoked through electrical stimulation of the neuropil and recorded using whole-cell patch-clamp electrophysiology of fate-mapped neurons at a holding potential of +40 mV. Recordings were performed at P8-12 and P17-21 from CTRL (NDNFCre-Ai14-NR1^+/+^) and NR1 KO (NDNFCre-Ai14-NR1^fl/fl^) L1 NDNF^+^ INs of vS1. The NMDA/AMPA response ratio was plotted between the two groups at both ages and statistically compared (P8-12 CTRL: 5 cells, P8-12 KO: 6 cells, P17-21 CTRL: 6 cells, P17-21 KO: 7 cells). **(e)** Two examples (above) with quantifications (below) of elicited action potentials within patched L1 interneurons, evoked through a train of 4 electrical stimulations of increasing frequencies (inter stimulus interval) (CTRL P17-21: 3 cells from 3 mice, KO P17-21: 4 cells from 2 mice). **(f)** L1 NDNF^+^ INs morphologically reconstructed. Sholl analysis was performed separately on axons and dendrites and statistical comparison was done for the overall average per distance step of 5 *μ*m (S.E.M. is shown around each point) (P8-12: 5 cells, P17-21: 5 cells). Statistics: (d), (e) and (f) two-tailed Mann-Whitney U-test (*p<0.05, **p<0.01,***p<0.001).

It has been shown that these interneurons display large NMDAR-mediated responses, which we hypothesized could impart them with the ability to integrate bilateral inputs (Chittajallu et al. 2020). To test this hypothesis we removed the mandatory subunit for NMDAR complex assembly (NR1) from these neurons by crossing the NDNFCre to the NR1*^fl^* mouse line, and ultimately obtained NDNFCre-NR1*^fl/fl^* offspring (denoted as KO). To assess the removal of the NR1 subunit we first performed a double fluorescent ISH against NDNF and NR1 mRNA. We saw that in contrast to control NDNF^+^ L1 cells, all of which expressed the NR1 subunit, no KO cells showed signal for NR1 at P21 (Figure 1c). To functionally validate the successful removal of the NMDAR, we electrophysiologically recorded *in vitro* evoked NMDAR-mediated responses from NDNF^+^ INs by stimulating locally, holding the membrane voltage (Vh) at +40 mV and blocking GABA*_A_*R/AMPAR/KainateR-mediated responses pharmacologically. The results showed that the NDNF^+^ NR1 KO cells indeed had significantly reduced evoked currents compared to CTRL (NDNFCre-NR1^+/+^), both at P8-12 and at P17-21 (Figure 1d). Note that even though we achieved a very strong reduction of the NMDAR-mediated currents, we did not obtain a full ablation (Figure 1d), in agreement with previous work in the hippocampus using a different ‘Cre’ driver mouse line (Chittajallu et al. 2017). To assess whether the reduction of NMDAR-mediated responses in KO NDNF^+^ INs would hinder their ability to integrate incoming inputs and subsequent action potential (AP) initiation, we performed a new set of whole-cell patch clamp recordings of CTRL and KO neurons in combination with stimulation of their incoming afferents, in current clamp mode. These experiments indeed showed that NMDARs contribute significantly to the capacity of NDNF^+^ INs to properly integrate incoming activity necessary to initiate AP firing (Figure 1e).

Morphological defects have previously been demonstrated to arise through embryonic removal of NMDAR from superficially located NGFC INs (García et al. 2015). Even though our manipulation is performed postnatally and past the developmental period responsible for these defects, ((García et al. 2015)), we directly tested the absence of a morphological phenotype in NDNF^+^ INs in our conditional genetic removal experiments. To do that, we labelled NDNF^+^ INs from both CTRL and KO mice with a fluorophore after filling the cells with biocytin and subsequently reconstructed their morphology at two developmental time points (Figure S1a). After tracing the cells’ neurites and assigning a dendritic or axonal identity to them, we measured the total length of the elements and found no difference in either the axon or the dendrites between CTRL and KO cells (Figure S1b). The same was true for the maximum distance that a cells’ neurites extend to (Figure S1b). Additionally, we applied Sholl analysis to assess the arborization of the neurites of CTRL and KO cells. The results showed that removal of the NR1 subunit resulted in no axonal changes at P8-12 or P17-21, and only very subtle and confined differences in the dendrites at P17-21, which could be accounted for by cell-to-cell variability (Figure S1c). These results indicate that the removal of NMDAR from NDNF^+^ INs in our experiments does not lead to a developmental morphological phenotype.

It is generally appreciated that the complexity of a cell’s neurites during development is a good indication for when it may functionally engage in the network activity. This would be especially true for NDNF^+^ INs, whose very elaborate axon is suggested to lead to a unusual spatiotemporal profile of GABA release and to the strong slow inhibitory effect they exert (Karayannis et al. 2010, Oláh et al. 2009, Szabadics et al. 2007). To assess when during development NDNF^+^ INs may begin participating in somatosensation, we analysed the morphology of CTRL neurons before and after P14, a time point right after mice increase their locomotor activity and start to bilaterally whisk to explore their environment. Our analysis showed an increase in the ramification of NDNF^+^ INs dendrites between P8-12 and P17-21, suggesting that these cells receive some information before whisking, which changes over time (Figure 1f top panel, Figure S1a). In contrast to the small dendritic change, there was a striking increase in the axonal length across the two time points, with these neurons having a significantly more elaborate axon at the late time point (Figure 1f bottom panel, Figure S1a). This indicates that it is only after the onset of bilateral whisking (P14) that these neurons would begin impacting the network with their uniquely slow output (Akhmetshina et al. 2016, Arakawa & Erzurumlu 2015). Additionally, the callosal fibres seem to also undergo an extensive expansion in arborisation around P11-12, granting support to the idea that the sensory flow from bilateral whisking is not yet mature before this time (Wang et al. 2007). As we aimed to investigate the role of L1 NDNF^+^ INs in coincidence detection and integration of incoming bilateral sensory information, we focused our subsequent experiments on P17-21 mice.

### L1 vS1 NDNF^+^ INs integrate bilateral whisker stimuli in an NMDA receptor-dependent manner to modulate the activity of underlying laminae

To assess if and how NDNF^+^ neurons integrate uni- and bi-lateral information to modulate the activity of underlying pyramidal neurons we performed acute *in vivo* silicon probe recordings within the vS1 barrels (Figure 2a). We then performed three sensory stimulation paradigms in each animal; the principal whisker, contralateral to the recording site was deflected once (denoted as ‘*contralateral*’), the corresponding ‘principal’ whisker at the ipsilateral side once (denoted as ‘*ipsilateral*’) or both simultaneously in a bilateral fashion (denoted as ‘*bilateral*’) (Figure 2b). First, we looked in CTRL mice, how multiunit activity (MUA) developed within the different laminae in response to all of the three paradigms. We functionally assigned layers to recording sites on the silicon probe based on current source density (CSD) analysis of the evoked responses (Reyes-Puerta et al. 2015, van der Bourg et al. 2017). We found that upon deflection of the principal contralateral whisker, cells of L1, L2/3 and L4 discharge a number of APs resulting in a characteristic largely bi-phasic temporal response with a fast, sharp and strong first peak, followed by a second more prolonged one (Figure 2c) (Vecchia et al. 2020). In contrast, we found that stimulation of the ‘principal’ whisker of the ipsilateral whisker pad lead to an overall delayed response that decayed slowly, in line with the longer travel time of the signal through the corpus callosum (Figure 2c). Combining the stimulation of both whiskers displayed a bi-phasic response that has a similar structure to the contralateral stimulation alone, but with a trend towards a shorter delayed phase of discharge in L2/3 (Figure 2c) (Palmer et al. 2012). Using the characteristic response profile of the evoked MUA within L4 we divided the responses into two windows used for analysis, an ‘*early*’ and a ‘*late*’, each corresponding to either of the two response ‘phases’ seen in the evoked activity (Figure 2a) (Yang et al. 2013). We then compared the CTRL *in vivo* MUA to the one recorded *in vivo* in the KO mice. We found that the information reaching the major thalamo-recipient L4 of the cortex did not display any significant change between the genotypes (Figure 2d). These functional data demonstrate that the activity carrying sensory-information via the lemniscal pathway is not altered at any relay station between the whisker and the barrel cortex (Bosman et al. 2011, Lu & Lin 1993), as expected based on the specificity of the genetic manipulation. In accordance with our *in vitro* electrophysiological data (Figure 1e) and hypothesis, we found that removing NMDARs from NDNF^+^ INs lead to a decrease in spiking activity recorded in L1, and that this was significant only upon the bilateral stimulus (Figure 2d). These data do not only demonstrate that the manipulation has functional validity *in vivo*, but importantly that the NDNF^+^ INs are dependent on NMDARs to integrate converging bilateral sensory information at distinct time scales (Figure 2d). In contrast to L1, when assessing the activity in CTRL vs. KO in L2/3 we surprisingly saw that there was a significant increase in the late phase of evoked MUA in the KO animals upon bilateral stimulation (Figure 2d), but no change in L5 (data not shown). These findings demonstrate not only that the NDNF^+^ population within L1 is responsible for integrating bilateral somatosensory signals in an NMDAR-dependent manner, but also that they use this sensory information to fine-tune the evoked activity in L2/3. *In vitro* Paired-cell recordings have shown that NGFCs have a high probability to be connected to L2/3 pyramidal cells and that they evoke strong and prolonged inhibitory postsynaptic potentials, partly through GABA*_B_* receptors (Schuman et al. 2019). Due to this and the fact that the NGFCs have distinct LS spiking properties, we hypothesized that the difference seen in the ‘late’ phase of L2/3 spiking could be accounted for by a change in the output of LS NGFC INs (Jiang et al. 2015, 2013, Schuman et al. 2019).

**Figure 2:**
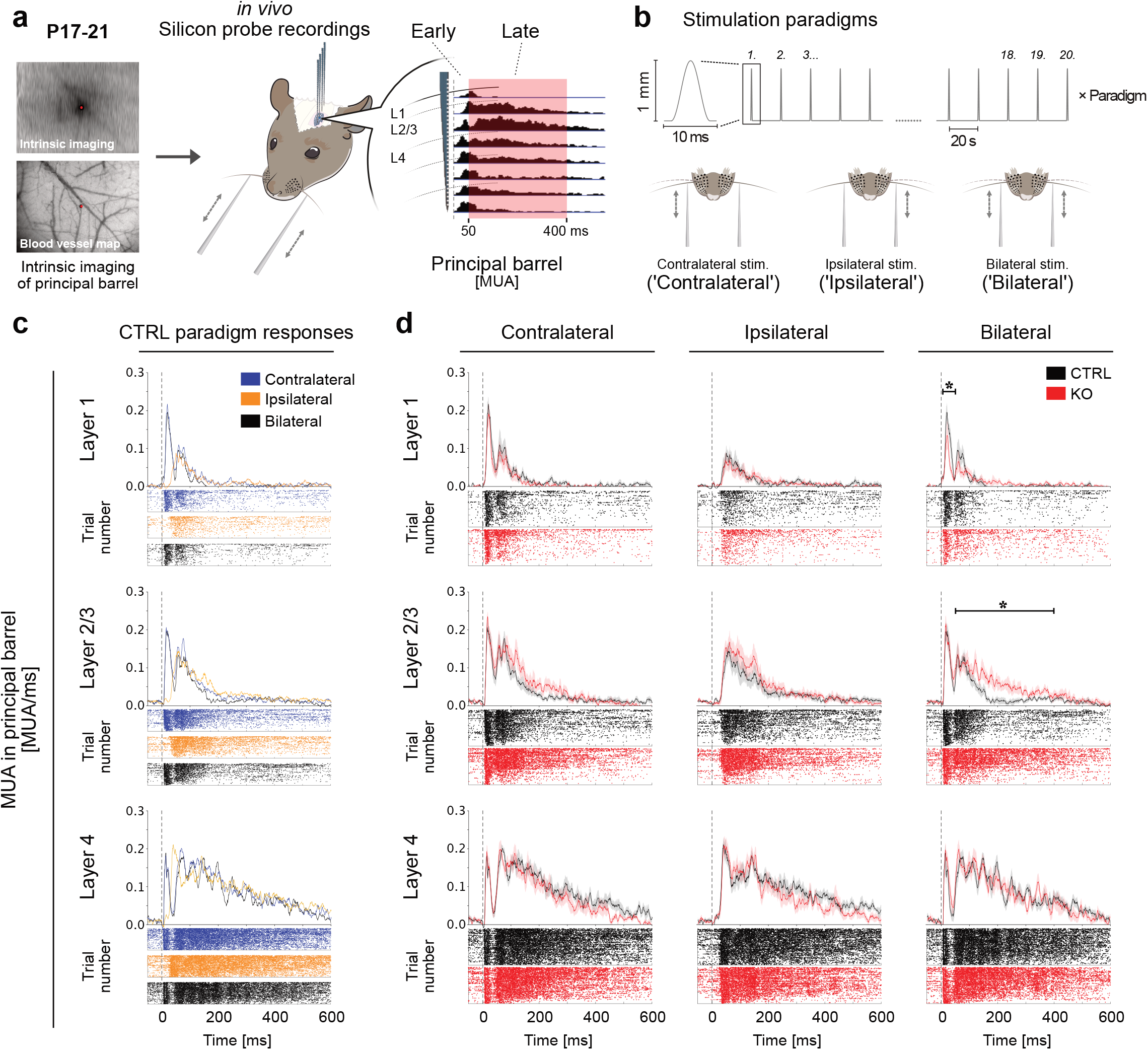
L1 vS1 NDNF^+^ INs integrate bilateral whisker stimuli in an NMDA receptor-dependent manner to modulate the activity of underlying laminae. **(a)** Silicon probes were implanted acutely *in vivo* in the principal column (C1 or C2) of CTRL or KO mice at P17-21 using intrinsic functional mapping in relation to the blood vessels. The laminae were functionally assigned to one of the eight recording sites of the silicon probe based on CSD analysis of the responses. Recorded MUA was separated into two temporal domains denoted ‘Early’ (5-50 ms) and ‘Late’ (50-400 ms) from the time of whisker deflection (0 ms), and statistically compared across groups for each of these domains. **(b)** Either the whisker contralateral to the recorded vS1 (‘Contralateral’), the whisker ipsilateral to this vS1 (‘Ipsilateral’) or both whiskers were displaced simultaneously (‘Bilateral’). Each whisker was displaced 20 times per stimulation paradigm with an 20s interval between each stimulus. **(c)** MUA over time for CTRL mice is plotted per lamina and per stimulation paradigm. The overall average is shown above and the raster plot representations of all activity from all stimulus presentations in all recording sessions is presented below. **(d)** MUA between CTRL vs. KO mice is plotted per lamina and per stimulation paradigm, with the overall average shown above (S.E.M. is shown as the shaded area around the average) and the raster plot representations of all activity from all stimulus presentations in all recording sessions is presented below. Statistical comparisons were done per temporal domain (CTRL: N=5 mice, n=9 principal columns; KO: N=5 mice, n=8 principal columns). Statistics: (d) two-tailed Mann-Whitney U-test (*p<0.05,**p<0.01,***p<0.001).

### L1 NDNF^+^ NGFCs modulate Layer 2/3 neuronal spiking activity in a GABA*_B_* receptor-dependent manner

To test if the L2/3 neuronal spiking effect seen in the KO mice is due to NGFC reduced output, we performed the same kind of *in vivo* silicon probe recordings as before in CTRL mice and assessed the activity in both L2/3 and L4, before vs. after the application of a GABA*_B_* antagonist (CGP 52432). The expectation was that abolishing activation of these receptors would mimic the effect we see in the KO mice. The first thing that we observed performing these experiments is that the application of the control or drug solution on the brain lead to a change in the spiking activity both in control and KO mice compared to the one recorded in the first set of experiments. Nevertheless, this acute experiment allowed to see that the MUA of L2/3 CTRL mice showed a clear change before vs. after the application of the GABA*_B_* antagonist on the cortical plate (Figure 3a), with increase of spiking in the late phase. In contrast, when performing the same experiment in the KO mice, we saw no difference within L2/3 MUA after the GABA*_B_* antagonist (Figure 3b). These results, together with the ones reported above, provide strong evidence that the GABA*_B_* receptors activated by L1 NGFCs are responsible for the bilateral stimulus-dependent attenuation of persistent activity in L2/3.

**Figure 3:**
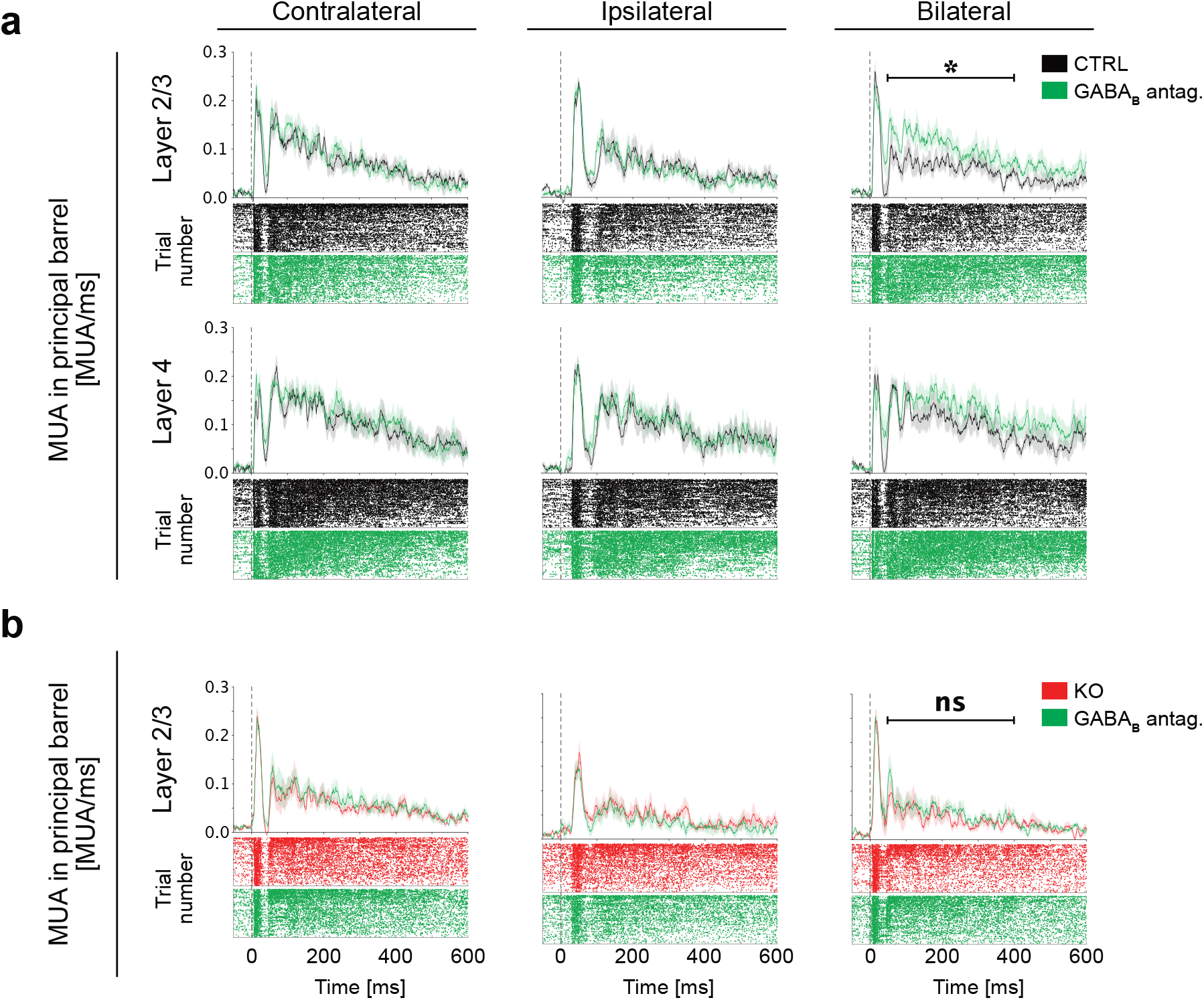
L1 NDNF^+^ NGFCs modulate underlying neuronal spiking activity in a GABA*_B_* receptor-dependent manner. **(a)** MUA between CTRL vs. CTRL+GABA*_B_* antagonist (CGP 52432) responses, plotted per lamina (horizontal panels) and stimulation paradigm (vertical panels), with the overall average shown above (S.E.M. is shown as the shaded area around the average) and the raster plot representations of all activity from all stimulus presentations in all principal columns and animals presented below. Statistical comparisons were done per temporal domain (CTRL: N=4 mice, n=7 principal columns; CTRL+GABA*_B_* antagonist (CGP 52432): N=4 mice, n=7 principal columns). **(b)** MUA between KO vs. KO+GABA*_B_* antagonist (CGP 52432) responses, plotted for L2/3 per stimulation paradigm, with the overall average shown above (S.E.MΓ. is shown as the shaded area around the average) and the raster plot representations of all activity from all stimulus presentations in all principal columns and animals presented below. Statistical comparisons were done per temporal domain (KO: N=5 mice, n=9 principal columns; KO+GABA*_B_* antagonist (CGP 52432): N=5 mice, n=9 principal columns). Statistics: (a) and (b) Wilcoxon signed-rank test for paired data (*p<0.05, **p<0.01, ***p<0.001).

### L1 NDNF^+^ NGFCs directly connect to and provide contralaterally-mediated feed-forward inhibition onto vS1 L2/3 pyramidal cells

To strengthen the functional data and assess the anatomical specificity of the circuit, we used retrograde monosynaptic pseudo-typed RV tracings to investigate the connectivity between L1 INs and L2/3 pyramidal cells in vS1. To achieve this, we performed *in utero* electroporation at embryonic day 15.5 (E15.5) to specifically express the TVA receptor and the glycoprotein ‘G’ to upper layer pyramidal cells. We subsequently injected the RV at P15 and sacrificed the mice at P21 (Figure 4a). Histological examination of slices from these brains showed that both superficial as well as deeper layer neurons connect to L2/3 pyramidal cells. A close look at L1 revealed scattered L1 IN labelling, with most cells found in the lower part of the layer, of which many expressed NPY, both indicative of NGFC INs (Figure 4b) (Schuman et al. 2019). These findings are in agreement with the higher probability of connection between L1 NGFCs and L2/3 pyramidal cells, compared to that between the latter and Canopy cells in an *In vitro* preparation (Schuman et al. 2019).

**Figure 4:**
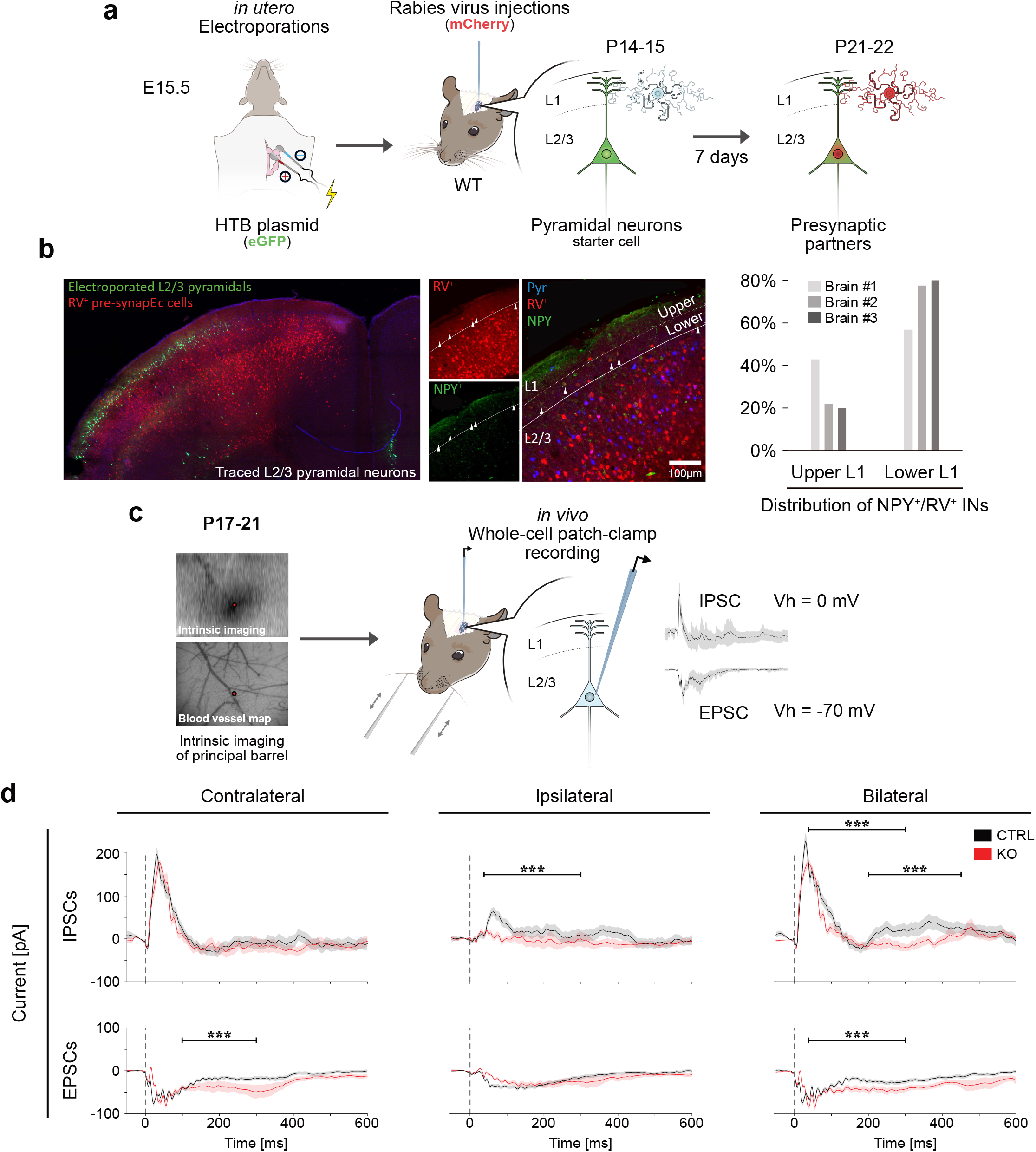
L1 NDNF^+^ NGFCs directly connect to and provide synaptic inhibition onto vS1 L2/3 pyramidal cells *in vivo*. **(a)** Schematic representation of the experimental paradigm for tracing monosynaptic connections onto L2/3 Pyramidal cells of vS1. *In utero* electroporation of an HTB-plasmid was performed in WT mice at E15.5. This was followed by RV injection into vS1 at P14-15 and the brain were extracted and analyzed at P21-22. **(b)** A representative photo of the mono-synaptically connected neurons of one hemisphere shown in red (left). A close-up of the injected hemisphere is shown. The pre-synaptic NPY^+^ partners within L1 of vS1 were separated into bottom or top half of L1 (middle image), and analyzed based on their position and NPY expression (right image and plot) (N=3 mice). **(c)** *In vivo* whole-cell voltage-clamp recordings from L2/3 neurons within the principal column (C1 or C2) of CTRL or KO mice at P17-21. Both EPSCs and IPSCs were recorded. These experiments were done in the presence of QX-314, while all three stimulation paradigms were presented. Recorded currents were separated into the three temporal domains; ‘Early’ (5-35 ms), ‘Late’ (35-100 ms) and ‘Slow’ (100-300 ms), from the time of whisker deflection (0 ms), and statistically compared based on these domains. **(d)** The grand average of IPSCs (top row) and EPSCs (bottom row) is shown together with the S.E.M. as the shaded area around (CTRL: N=6 mice, n=10 neurons; KO: N=7 mice, n=7 neurons). Statistics: (d) two-tailed Mann-Whitney (*p<0.05, **p<0.01,***p<0.001).

To reveal in detail how the NGFCs of L1 integrate bilateral information to modulate underlying pyramidal neurons, we turned to *in vivo* intracellular recordings. *In vivo* whole-cell voltage-clamp recordings from L2/3 pyramidal cells of either CTRL or KO mice allowed us to dissect how incoming inhibitory post synaptic currents (IPSCs) and subsequently excitatory post synaptic currents (EPSCs) were altered due to the L1 NDNF^+^ manipulation (Figure 4c). We first looked at the whisker-evoked IPSCs (Vh = 0 mV) arriving to the recorded cells after the three stimulation paradigms. In accordance with our MUA findings and hypothesis, the overall evoked incoming inhibition was significantly reduced within the L2/3 neurons when either the ipsilateral or the bilateral paradigm was used (Figure 4d, top row). Interestingly, a second late phase of incoming inhibition was visible in both paradigms, but was much more clear when the bilateral stimulation paradigm was applied. Even though both the fast and the slow IPSC were reduced in the KO mice after the bilateral stimulation, the effect on the second peak was more pronounced (Figure 4d, top row, right). Furthermore, these recordings also revealed a very intriguing and unexpected finding with the incoming inhibition evoked through the ipsilateral stimulation paradigm almost completely abolished after removal of NMDARs from L1 NDNF^+^ Ins. This suggests that the vast majority (if not all) of feed-forward inhibition onto L2/3 pyramidal cells coming from the opposite hemisphere is provided by L1 NDNF^+^ INs (Figure 4d, top row, middle). The time course of the late inhibitory synaptic current in the CTRL bilateral stimulation paradigm is suggestive of GABA*_B_*R activation and overlaps well with that of the ‘late’ phase of the L2/3 MUA. The prolonged incoming excitation and the lack of incoming inhibition from around 200 ms onwards within KO neurons overall matches the increase in the MUA seen in the silicon probe recordings of bilateral L2/3. This demonstrates that at the single cell level, the increase in the late phase of bilateral L2/3 activity in the KO is due to a reduction in neuronal inhibition coming from L1 NDNF^+^ cells, but also to some extent through a change in excitation, most likely due to the lack of inhibitory control onto the surrounding presynaptic pyramidal neurons. This was apparent by looking at the isolated EPSCs (Vh = −70 mV) onto pyramidal cells in CTRL vs. KO mice after implementing the three whisker stimulation paradigms (Figure 4d, bottom row). The main difference between the overall incoming currents within the KO mice was seen in the late phase of the contralateral, as well as the bilateral paradigm, with a temporal increase in incoming excitation compared to that found in CTRL neurons (Figure 4d, bottom row). The timelines seem to match well with the extended period of registered MUA within L2/3 when comparing between the same stimulation paradigms. It is interesting to note that while differences were seen in EPSCs evoked by the contralateral and bilateral paradigms, no apparent change was observed when performing the ipsilateral stimulation paradigm (Figure 4d, bottom row).

Taken together, our data demonstrate that there is a specialized feed-forward inhibitory circuit controlling L2/3 pyramidal cells in somatosensory cortex, particularly upon bilateral sensory stimulus presentation. This stream of feed-forward inhibition results in a delayed and long-lasting inhibitory effect on the discharge of L2/3 pyramidal cells, via NMDAR-dependent engagement of L1 NGFC INs.

## Discussion

In this study we address how bilateral information is processed in the somatosensory cortical area of the mouse that receives whisker-related information, and how L1 NDNF^+^ inhibitory cells regulate this information. Using silicon probes we record the activity of neurons of the barrel cortex after contralateral, ipsilateral or bilateral principal whisker deflection. Firstly, we functionally show the preferential homotopic information transfer among the ‘same’ corresponding barrels across the midline. In accordance with previous work, we find that upon deflection of the principal contralateral whisker, pyramidal cells of L2/3 discharge APs in a characteristic bi-phasic temporal manner with a fast, sharp and strong first peak, followed by a second smaller but more prolonged peak (Vecchia et al. 2020). In contrast, we find that stimulation of the respective principal whisker of the ipsilateral whisker pad has a delayed onset, in line with the longer travel time of the signal through the corpus callosum. Combining the stimulation of both whiskers displays a bi-phasic response that has an overall similar structure to the contralateral stimulation alone. Our results also show that inhibition is impacting the late spiking activity of L2/3 cells particularly upon bilateral stimulation via not only a GABA*_A_*R, but also in a GABA*_B_*R-dependent manner. Based on our rabies tracing experiments onto L1 NDNF^+^ INs and L2/3 pyramidal cells, as well as the conditional genetic manipulation, we find that these GABA*_B_*R are activated by NGFCs. This capacity of L1 NGFCs to be activated by bilateral stimuli is not only conferred by the bilateral inputs they receive, but also by a molecular program that is activated in them that imbues them with large NMDAR-mediated responses. The phenomenon we have uncovered on the role of these cells in bilateral processing seems not to be specific only to the somatosensory cortex, but perhaps to all sensory cortices since the long-range connections have been shown to exist in primary auditory cortex as well (Abs et al. 2018). Our findings are also consistent with previous work showing that NGFCs of L1 have a high probability of connection to L2/3 pyramidal cells (Schuman et al. 2019), identified by paired whole cell recordings. Our work makes the surprising finding though that these inhibitory connections provide a strong delayed inhibitory control only over the late L2/3 pyramidal cell-spiking phase upon bilateral whisker stimulation, but not that of L5 cells. Interestingly, this L2/3 late spiking phase has been shown to be key for sensory perception, suggesting that NDNF^+^ INs have a key role to play in the perception of bilateral tactile stimuli (Libet et al. 1967, Sachidhanandam et al. 2013). Previous work has also found that delayed slow IPSPs (sensitive to GABA*_B_*R blockade) onto L5 pyramidal cells, obtained after callosal stimulation *in vitro*, were not abolished by blocking AMPARs/KainateRs in Mg^2+^-free solution. Nevertheless, these slow IPSPs were sensitive to both GABA*_B_*R as well as NMDAR blockade, suggesting the engagement of a strong NMDAR-dependency in slow feed-forward inhibition (FFI) (Kawaguchi 1992). In contrast, by looking at early occurring inhibitory responses in the same experiment and performing similar pharmacological manipulations, the author concluded that NMDARs seem to be less involved in the excitation of neurons mediating fast GABA*_B_*R-dependent FFI onto L5 pyramidal cells (Kawaguchi 1992). Intriguingly, our observations of spiking activity in L5 being moderately decreased after removal of NMDARs from L1 NDNF^+^ neurons (data not shown), suggest that bilateral stimuli may exert disinhibition onto the majority of L5 pyramidal cells via L1 NDNF^+^ cells. Even though this result may appear to be in disagreement with previous research looking at the spiking activity of individual L5 pyramidal cells *in vivo* and *in vitro* coming from the contralateral hemisphere, (Anastasiades et al. 2018, Palmer et al. 2012), we also find that application of a GABA*_B_*R antagonist in the cortex of WT animals suppresses the late activity of L5 pyramidal cells (data not shown) (Anastasiades et al. 2018, Palmer et al. 2012). These results therefore suggest that L1 NDNF^+^ INs provide strong FFI onto L2/3 pyramidal cells predominantly upon bilateral stimuli, but not onto L5, which would be provided by a different set of cells (Niquille et al. 2018) (Silberberg & Markram 2007, Urban-Ciecko et al. 2015). Our intracellular recordings of L2/3 cells showing almost complete loss of IPSCs in the KO upon ipsilateral stimulation further propose that L1 NGFCs are the major providers of FFI onto L2/3 pyramidal cells coming from the contralateral hemisphere. Interestingly, the effect of NMDAR removal on the late phase of the L2/3 spiking activity is not only in line with the slow synaptic output of NGFCs, but also with their late spiking characteristics. Previous work using intracellular recordings in cat visual cortex has speculated that inhibitory responses evoked by corpus callosum or thalamic LGN stimulation onto upper- and lower-layer pyramidal cells are produced by the same inhibitory cell, suggesting convergence of these excitatory inputs onto a single IN type. Based on our rabies experiments and our silicon probe findings we would propose NGFCs to play this role.

Our results also suggest that the ‘remaining’ NDNF^+^ Canopy cells might in contrast primarily target inhibitory cells that in turn provide inhibition to L5 pyramidal cells, in line with what has also been reported by Toyama and Matsunami when stimulating thalamocortical and callosal afferents with slight delays (Toyama & Matsunami 1976). More recent research has shown that some L1 5HT3aR^+^ INs, to which NDNF^+^ cells belong, provide inhibitory inputs to L4 PV^+^ cells in the auditory cortex, which are important for sound-evoked plasticity (Takesian et al. 2018). Further recent findings in the mouse visual cortex have also shown that L1 NDNF^+^ cells provide disinhibitory somatic control over L5 pyramidal cells spiking activity via PV^+^ cells (Malina et al. 2021). In accordance with these results, NDNF^+^ cells of the prefrontal cortex (PFC) were shown to provide stronger inhibition onto superficial PV^+^ neurons compared to SST^+^ INs (Anastasiades et al. 2020). In conclusion, the work presented herein, together with published work on other parts of the cortex suggest that different streams of inhibition are activated by ipsilateral stimuli onto distinct layers of the cortex. The L2/3 cells display typical FFI coming from the contralateral cortex and mediated via L1 NGFCs, whereas L5 cells receive a combination of FFI and disinhibition. The two latter ones could be provided at distinct timescales on different subcellular compartments of L5 cells or even on two distinct L5 pyramidal cells that are controlled in a differential manner via different IN types when processing bilateral information. These multiple FFI streams have important implications for sensory coding and action planning, as depending on whether the pyramidal cells are Intratelencephalic or Pyramidal tract neurons, they would not only display unique dendritic integration properties, but importantly would activate or suppress unique downstream targets (Takahashi et al. 2020).

## Materials and Methods

### Animals

All animal experiments were approved by the Cantonal Veterinary Office Zürich and the University of Zürich and were approved by the veterinary office of the canton of Zürich. Animals were housed in a 12-h reverse dark-light cycle (7am to 7pm dark) at 24°C at variable humidity. Animal lines used in this study are: Ndnf-IRES2-dgCre-D (B6.Cg-Ndnf^*tm*1.1(*folA/cre)Hze*^/J) (Tasic et al. 2016), fNR1 (B6.129S4-Grin1^*tm*2*Stl*^/J) (Tsien et al. 1996), B6/J-Rj (C57BL/6JRj), HTB (B6-Gt(ROSA)26Sor^tm1(*CAG–neo,–HTB*)*Fhg*)^ (Li et al. 2013) and tdTomato/Ai14 (B6;129S6-Gt(ROSA)26Sor^*tm*14(*CAG–tdTomato*)*Hze*^/J) (Madisen et al. 2010). Because the Ndnf-IRES2-dgCre-D was originally designed to be inducible through trimethoprim (TMP) injections, we injected all mice postnatally with TMP of 1.25 mg/g body weight (Sando et al. 2013).

### Fluorescent RNA in situ hybridization

Postnatal mice were perfused with 5ml of phosphate saline buffer 1 × (PBS) and 10ml of paraformaldehyde 4% in phosphate buffer 0.1M (PFA 4%); brains were dissected and then post-fixed in PFA 4% for 2 hours. Fixed tissue was cryoprotected with sucrose 30% in PBS overnight and then embedded in OCT and stored at −80°C. Brains were sectioned (25 *μ*m) with a cryostat on slides (Superfrost Plus, Thermo scientific) and stored at −80°C. Slides were defrosted, air dried for 1 hour, fixed in formaldehyde 4% in PBS for 10 min, washed twice in PBS for 5 min, fixed in 1.5% H2O2 in methanol for 15 min, washed twice in PBS for 5 min, treated with 0.2M HCl for 8 min, washed twice in PBS for 5 min, digested with Proteinase K (10 g/ml in PBS) for 5 min, washed once in PBS for 5 min, incubated in acetylation solution (for 200 ml of solution: 2.66 ml triethanolamine >99.5%, 0.32 ml HCl 37%, 0.5 ml acetic anhydride 98%) for 10 min, washed three times in PBS for 5 min. Slides were then placed in a humid chamber and covered with hybridization solution (formamide 50%, SSC 5 ×, yeast tRNA 0.1 mg/ml, Denhardt’s solution 1 ×, salmon sperm 0.1 mg/ml) for at least 2 hours and then hybridized with RNA probes overnight at 65°C. Prior incubation, RNA probes were diluted 1ng/*μ*l in hybridization solution, heated at 80°C for 5 min and then immediately transferred on wet ice for 3 min. Slides were washed once in SSC 5× for 5 min at 65^°^C, twice in formamide 50%, SSC 2× for 30 min at 65°C, once in SSC 2× for 15 min at 37°C, once in SSC 0.1 × for 15 min at 37°C and once in TN buffer (Tris-HCl pH 7.5 0.1M and NaCl 0.15M). Slide were moved to a humid chamber and blocked with blocking solution (MAB 1×, blocking reagent (Roche) 2%, Tween-20 0.3%) for 1 hour and finally incubated overnight at 4°C with anti-Dig-POD antibody (Roche) diluted 1:500 in blocking solution. Slides were washed in TNT buffer (TN buffer, Tween-20 0.05%) three times for 5 min and once for 1.5 hours. Dig-RNA probes were developed with Cy5-tyramide 1:100 in amplification reagent (PerkinElmer) for 20 min. Slides were washed in TNT buffer three times for 5 min, incubated in 3% H2O2 in TN buffer for 2 hours, washed in TNT buffer three times for 5 min, placed in a humid chamber, blocked with blocking solution for 1 hour and finally incubated overnight at 4^°^C with anti-Fluorescein-POD antibody (Roche) diluted 1:500 in blocking solution. Slides were washed in TNT buffer three times for 5 min and once for 1.5 hours. Fluorescein-RNA probes were developed with Fluorescein-tyramide 1:100 in amplification reagent (PerkinElmer) for 20 min. Slides were washed in TNT buffer three times for 5 min, washed once in TN buffer for 5 min and mounted with Fluoromount-GTM mounting medium with DAPI (Invitrogen).

### RNA probes

Genes of interest were cloned from mouse postnatal brain cDNA using the following set of primers; Gad1 Forward: TGTGCCCAAACTGGTCCT, Gad1 Reverse: TAATACGACTCACTATAGGGTGGCCGATGATTCTGGTT,NR1 Forward: ACCTTGTGGCAGATGGCAAG, NR1 reverse: TAATACGACTCACTATAGGGATGGCCTCAGCTGCACTC, Ndnf Forward: GCGATGCACCTTTGGAGT, Ndnf Reverse: TAATACGACTCACTATAGGGGACAGAAGCAGCCTCCCA. All primers have been provided by the Allen Brain Institute developing brain atlas (Atlas. 2004a), except for the “NR1 Forward”. This primer was designed to specifically target the sequence flanked by the loxP sites in the NR1 conditional KO mouse. RNA probes have been transcribed using a T7 polymerase (Roche) with either Dig-NTPs or Fluorescein-NTPs (Roche) and purified with Post Reaction Clean-Up Columns (Sigma).

### Slice Preparation

NR1 Control (NDNFCre-Ai14-NR1+/+) and NR1 KO mice (NDNFCre-Ai14-NR1fl/fl), between P17 and P21, were anesthetized by isofluorane and decapitated, and their brains were rapidly transferred to ice-cold dissecting aCSF containing (mM): 125 NaCl, 2.5 KCl, 25 NaHCO_3_, 1.25 NaH_2_PO_4_, 10 MgCl_2_, 0.5 CaCl_2_ and 22 glucose, saturated with 95% O2 and 5% CO2. *Ex-vivo* acute coronal brain slices (300 *μ*m thick) were cut by using a Vibrotome (VT 1200S, Leica), then transferred to a recovery bath filled with aCSF containing (mM): 125 NaCl, 2.5 KCl, 25 NaHCO_3_, 1.25 NaH_2_PO_4_, 1 MgCl_2_, 2 CaCl_2_ and 22 glucose continuously aerated with 95% O2 and 5% CO2. Slices were kept at room temperature and were continuously perfused with aCFS at a rate of 2-3 ml/min at 30°C during experiments.

### Patch-clamp recordings

Patch-clamp recordings in whole-cell configuration were performed on layer 1 (L1) interneurons (INs) in whisker primary somatosensory cortex. Borosilicate patch pipettes (3–4 MΩ) filled with a solution containing (in mM): 130 KMeSO4, 5 KCl, 5 NaCl, 10 HEPES, 0.1 EGTA, 2 MgCl2, 0.05 CaCl2, 2 Na2-ATP and 0.4 Na3-GTP (pH 7.2-7.3, 280-290 mOsm/kg) were used for the recordings. NDNF^+^ INs in L1 of whisker primary somatosensory cortex were visualized under infrared-differential interference contrast (DIC) and identified through fluorescence. Current-Clamp experiments were performed, and post-synaptic potentials (PSPs) were evoked by using a monopolar stimulation through a glass pipette positioned in Layer 2/3 (L2/3) connected to a constant-current isolation unit and acquired every 10 seconds. Stimulation parameters (0.43 mA to 0.72 mA) were set by evoking an averaged PSP responses of 11 mV to 14 mV over 10 evoked PSPs. After setting the stimulation parameters, sets of trains of 20Hz, 40Hz, 50Hz, 100Hz and 200Hz, were delivered 10 seconds apart. Each stimulation set was composed by trains of 4 stimuli, repeated 10 times, and acquired every 10 seconds. Patched INs were clamped at −70mV all over the recordings. Data were acquired using a Axopatch 200B amplifier controlled by pClamp software (v10.7.0.3, Molecular Devices 2016), filtered at 10 kHz and sampled at 10 kHz (Digidata 1440A, Molecular Device). All data are reported without corrections for liquid junction potentials. Data where the input resistance (Rinp) changed >20% have been excluded.

### Electroporations, Viral Injections and Monosynaptic Rabies Tracings

For the NDNF-tracing experiments, electroporation of NDNFCre mice pups at P2 with cre-dependent Dlx5/6-HTB plasmid (Dlx5/6-loxp-STOP-loxp-HTB) were conducted (De la Rossa & Jabaudon 2015), allowing a sparse, but strong expression of the TVA receptor and the glycoprotein “G” in NDNF^+^ INs of L1. The macro vibrissae barrel field within pups was subsequently injected with ASLV-A envelope glycoprotein (EnvA)-pseudotyped, glycoprotein-deleted rabies virus SADG-mCherry(EnvA) (Wickersham et al. 2007) at P14. 300 nl of the RV were topically applied (15 nl/s) on top of the barrel field of the S1. Stereotaxic vS1 coordinates (mm) at P14; AP: ~1.6, ML: ~3.0. The application was carried out using a glass micropipette attached to a Nanolitre 2010 pressure injection apparatus (World Precision Instruments). After the surgeries, the animals were monitored and returned to their home-cage for 10 days until P24, to allow for adequate viral expression.

For the L2/3 pyramidal-tracing experiments, C57BL/6 mice (denoted WT) embryos were co-electroporated with pCAG-HTB and pCAG-GFP plasmids at E15.5 in order to target the L2/3 pyramidal neurons. At P14-15 the electroporated pups were injected in the macro vibrissae barrel field with 200 nl (15 nl/s) of glycoprotein-deleted rabies virus SADG-mCherry (EnvA) (Wickersham et al. 2007). The RV was topically applied on top of the barrel field of the primary somatosensory cortex. The application was carried out using a glass micropipette attached to a Nanolitre 2010 pressure injection apparatus (World Precision Instruments). After the surgery, the animals were monitored and returned to their home-cage for 7 days until P21, to allow for adequate viral expression. At P21 the injected mice were perfused with 10 ml PBS, subsequently 15 ml PFA and their brains were post-fixed for 2 hours. The brains were cut with the vibratome apparatus (Leica) in 100 **μ**m thick slices and then stained with anti-rabbit NPY antibody (1:500, Immunostar).

### Confocal Imaging and Image Processing of Fate-Mapped NDNF and Pseudo-typed Rabies Traced Tissue

For NDNF fate-mapped (NDNFCre-Ai14/tdTomato) tissue, all images were taken from the cortical plate of the vS1 or contralateral vS1. For each of the processed brains stacks of 10 images were taken. For imaging a 20× oil-immersion objective (HC PL APO CS2, 20×/0.75 IMM) was used and stacks were taken of 0.5 *μ*m distance.

For in situ hybridization staining, 20× glycerol immersion objective (HC PL APO CS2 20×/0.75 IMM) was used and stacks were taken of 1 *μ*m distance.

For NDNF-tracing data, all images from the injection sites were taken with the FV100 Olympus confocal microscope using 20× objective in order to count the RV-infected cells (expressing mCherry) in L1. The percentage of rabies infected cells that also expressed NPY was calculated. The position of these cells in L1 was estimated by dividing L1 into two equal domains (an upper and a lower).

### Acute Slice Electrophysiology

Whole-cell patch-clamp electrophysiological recordings were performed on either NDNFCre-tdTomato or NDNFCre-fNR1-tdTomato cells in L1 of the vS1, in acute slices prepared from P8-12 or P17-21. Animals were anesthetized, decapitated, the brain extracted and transferred to 4°C physiological Ringer’s solution (aCSF), of the following composition (mM): 125 NaCl, 2.5 KCl, 25 NaHCO3, 1.25 NaH2PO4, 1 MgCl2, 2 CaCl2 and 20 glucose. The brain was then glued to a stage and cut into 300 *μ*m-thick coronal slices using a vibratome (VT 1200S, Leica). The slices recovered in room temperature aCSF for 30 min before recording. The slices were then placed in the recording chamber of an upright microscope (Axioscope 2 FS, Zeiss) and superfused with 32°C oxygenated (95% O2 and 5% CO2) aCSF at a rate of 2-3 ml/min. The microscope was equipped with immersion differential interference contrast (DIA) and the following objectives were used to visualize the cells (10×/0.3, Olympus and 40×/0.8, Zeiss). A CMOS camera (optiMOS, QImaging) was attached to the scope to visualize the slice and cells through a computer screen. A white-light source (HAL 100, Zeiss) and a LED based excitation source (Polygon400, Mightex Systems) in combination with a tdTomato filter set (set 43 HE, Zeiss, Excitation 550/25, Emission 605/70) were used to locate the fluorescent INs. Patch pipettes were pulled from borosilicate glass capillaries (1.5 OD × 0.86 ID × 75 L mm, Harvard Apparatus) at a resistance of 3-5 MΩ. For recordings of electrically evoked currents, Clampex was used (v10.7.0.3, Molecular Devices 2016). The recording pipettes were filled with a solution containing the following (mM): 135 potassium D-gluconate, 4 NaCl, 0.3 Na-GTP, 5 Mg-ATP, 12 phosphocreatine-di-tris, 10 HEPES, 0.0001 CaCl2 (pH 7.25, mOsm 290). In some cases 3 mg/ml biocytin (Tocris) was added in the recording solution. Electrical stimulator was positioned within L2/3 roughly 150 *μ*m below the patched L1 IN. Pharmacology was added to the aCSF superfuseing the tissue. Recordings were done in voltage clamp (Vh = +40 mV), while 20 **μ**M Gabazine (SR 95531 hydrobromide, Cat. Nr. 1262, Tocris) was added to allow us to measure evoked excitatory currents without contamination of inhibitory currents. Gabazine and 20 *μ*M CNQX (Cat. Nr. 1045, Tocris) was added to isolate NMDAR-mediated currents. Gabazine, CNQX and 50 *μ*M AP5 (Cat. Nr. 0106, Tocris) was added for validity of the measured NMDAR currents. All pharmacology was washed in for at least 10 min before recording. Currents were evoked within the patched cells through the electrical stimulator and the AMPAR-mediated component was calculated by subtracting the NMDAR component from the mixed AMPAR/NMDAR response. Access resistance was constantly monitored to ensure the quality and stability of the recording. The recorded data were accepted only if the initial series resistance was less than or equal to 25 MΩ and did not change by more than 20% throughout the recording period. Compensation was made for the pipette and cell capacitance. The recorded data was sampled at 10 kHz.

### Morphological Reconstructions and Structural Analysis

NDNFCre-tdTomato or NDNFCre-fNR1-tdTomato brain tissue containing biocytin filled NDNF^+^ L1 INs were cut in 150 *μ*m coronal sections (VT 1200S, Leica). Tissue was washed 3 × 10 min in PBS. A mix of streptavidin conjugated alexa fluor 488 (1:250) (Cat. No. S32354, Invitrogen), Triton (0.25%) and PBS was added to the free-floating slices, incubating for 48 hours at gentle shake in 4°C. Slices were rinsed for 3 × 15 min in PBS and cover-slipped with DAPI Fluoromount-G (Cat. Nr. 0100-20, SouthernBiotech). Tissue slices was imaged using a Confocal Microscope (Olympus FV1000 with Fluoview software). Z-stacks of 100-200 images were taken of INs in L1 of the vS1. Stacks were taken with a 60× silicone oil-immersion objective, stepsize of 0.5 *μ*m and resolution of 1600 × 1600 px. Neurons were traced using the Simple Neurite Tracer tool (Fiji, ImageJ) and axons as well as dendrites were separated. A continues sholl-analysis was applied with Fiji, values extracted and CTRL across age or CTRL vs. KO was plotted using Python Matplotlib. Statistical comparison was made between CTRL vs. KO per set distance of 5 *μ*m.

### Intrinsic Optical Imaging

The principal whisker-related barrel column was identified using optical imaging of intrinsic signals. The cortical surface was visualized through the intact bone by surface application of normal Ringer’s solution and a glass coverslip placed on top. The skull surface above the barrel cortex was thoroughly cleaned and left intact. Reference images of the cortical blood vessel pattern were visualized by a 546-nm LED to enhance contrast. Functional maps of the target macro vibrissae barrel column were obtained by shining red light (630 nm LED) on the cortical surface while stimulating the barrels corresponding whisker with a miniature solenoid actuator (described below in detail). Reflectance images were collected through a 4× objective with a CCD camera (Scientifica SciCam Pro; 16-bit; binned at 2-by-2-pixels, 688 × 512 binned pixels at 20 fps). Functional intrinsic signal images were computed as fractional reflectance changes relative to the pre-stimulus average (average of 15 trials). The intrinsic signal image obtained for the C1 or C2 barrel column was then mapped to the blood vessel reference image and used to guide the location of the craniotomy for both the *in vivo* multi-electrode recording and the *in vivo* patch-clamp electrophysiology experiments.

### Animal Surgery and Preparation for *in Vivo* Electrophysiology

In total we used 15 NDNFCre mice and 16 NDNFCre-fNR1 at P17-21 for the *in vivo* multi-electrode recordings and the *in vivo* patch-clamp experiment. Mice were sedated with chlorprothixene (0.1 g/kg, intraperitoneal (i.p.); Sigma-Aldrich Chemie GmbH, Buchs, Switzerland) and lightly anesthetized with urethane throughout the experiment (0.25–0.5 g/kg, i.p.). Atropine (0.3 mg/kg; Sigma-Aldrich Chemie GmbH, Buchs, Switzerland) and dexamethasone (2 mg/kg; aniMedica GmbH, Senden-Bösensell, Germany) were injected subcutaneously (s.c.) 30mins after induction of anesthesia to reduce salival secretion and prevent edema. Body temperature was kept at 37° C using a heating pad. Hydration levels were maintained by s.c. injections of Ringer-lactate (Fresenius Freeflex; Fresenius Kabi AG, Oberdorf, Switzerland) and depth of anesthesia was checked throughout the experiment by pinching the forepaw and checking for reflex. To stabilize the animal, a custom-built head plate was placed on the skull over the left hemisphere using first glue (Loctite 401, Henkel & Cie. AG.) and then dental cement (Paladur, Heraeus Kulzer GmbH Hanau, Germany; Caulk Grip Cement for electrophysiology). Using a sharp 26 gauge needle (26G × 1/2” Sterican, B. Braun) a small cranial window was created above the center of the mapped barrel column(s). Cranial window was about 2.0 × 2.0 mm^2^ in size for the multi-electrode recordings and 0.15 × 0.15 mm^2^ for patch-clamp electrophysiology experiments. Exposed dura was superfused with Ringer’s solution (Ringer’s, B. Braun). Care was taken not to damage the dura or surface blood vessels.

### *In Vivo* Multi-Electrode Silicon Probe Recordings

Neural activity was recorded with an 8 shank 64-channel ‘silicon probe’, inserted perpendicularly into vS1 (A8 × 8-Edge-5mm-100-200-177, NeuroNexus Technologies). Each of the shanks (5 mm long, of which the 750 *μ*m tip contained the recording sites) contained 8 recording sites (177 *μ*m^2^ surface area per recording site) spaced 100 *μ*m apart. Distance between each shank was 200 *μ*m. Insertion of the probe was guided by intrinsic optical imaging. For selected animals, the probe insertion points were marked by impregnating the probes with DiI (Cat. No. D282, Molecular Probes) before insertion. A silver wire placed over the cerebellum served as a ground electrode. All data were continuously digitized at 20 kHz and stored for offline analysis using a 64-channel extracellular recording system (SC2 × 32 and a 100-1-500 SN. 127, Multi Channel Systems MCS Gmbh) and MC_RACK software (Multi Channel Systems MCS Gmbh). One hour following electrode insertion the recordings were started. The total duration of the multi-electrode recordings varied between 3 and 7 h. After the recordings, mice were euthanized, perfused with PBS and the brains was put in 4% PFA overnight.

For a set of experiments pharmacology was used, either in NDNFCre or NDNFCre-fNR1 mice. Experiments were carried out in similar manner as described above with the difference that the whole experiment was divided in two parts, first part without pharmacology and second part with pharmacology. Before recording of the first part normal Ringer’s solution (Ringer’s, B. Braun) was applied to the top of the craniotomy for 30 min before recording. For the second part, a mixture of Ringer’s and 1 *μ*M of the GABA*_B_* antagonist CGP 52432 (Cat Nr. 1246, Tocris) was topically applied and recordings were initiated after 30 min (Iurilli et al. 2012). When washout was carried out, Ringer’s solution was applied and recordings were started 60 min later.

### *In Vivo* Whole-Cell Patch-Clamp Recordings

Patch pipettes were pulled from borosilicate glass capillaries (1.5 OD × 0.86 ID × 150 L mm, Harvard Apparatus) at a resistance of 5-7 MΩ. Pipettes were filled with internal solution (mM): 135 potassium D-gluconate, 4 NaCl, 0.3 Na-GTP, 5 Mg-ATP, 12 phosphocreatine-di-tris, 10 HEPES, 0.0001 CaCl2 (pH 7.25, mOsm 290), and 3.5 mg/ml QX-314 Bromide was added (Cat No. 1014, Tocris). Signal was amplified (Multiclamp 700B, Molecular Devices), digitally converted (Axon Digidata 1550A, Molecular Devices) and recorded at 10 kHz through Clampex (v10.7.0.3, Molecular Devices 2016), while a silver pellet submerged within the solution covering the craniotomy served as a ground. An opening of the dura was made with the first pipette. High positive pressure (~100 mbar) was applied outside the brain and reduced (~30 mbar) after passing the pia mater. Pipette tip was first submerged into the brain until wanted depth and then subsequently submerged in a stepwise fashion (2 *μ*m/step) until neuronal contact. If contact was successful, gigaseal and whole-cell configuration was established. The patched cell was acclimatized to the intracellular solution for 3 min before recording. Recording of whisker evoked currents were done in voltage clamp at either Vh = −70 mV (to isolate EPSCs) or Vh = 0 mV (to isolate IPSCs). After the recordings, mice were euthanized, perfused with PBS and extracted brains postfixed overnight in 4% PFA.

### Whisker Evoked Stimulation

After intrinsic imaging of a corresponding macro vibrissae, remaining whiskers were trimmed. A single whisker pair was deflected 1 mm from the whisker pad using a miniature solenoid actuator adapted from Krupa et al (Krupa et al. 2001), driven by a stimulus generator (STG4002-160 *μ*A, Multi Channel Systems MCS Gmbh) connected with an in-house 10 × voltage amplifier. Stimulator generated for 10 ms a 1000 *μ*m displacement in the anterio-posterior direction. Recordings were carried out while either; the whisker contralateral to the recording site, the whisker ipsilateral to the recording site or both whiskers of a corresponding pair were deflected in a bilateral manner. For the *in vivo* patch-clamp experiments, the 3 whisker stimulation paradigms were presented in a repeated sequence of ‘*Contra*’, ‘*Ipsi*’ and ‘*Bilat*’ stimulation, until 20 sequences had been presented per recording. For the *in vivo* multi electrode recordings, the three stimulation paradigms were presented separately in a pseudorandom manner, until 20 repetitions had been reached per paradigm. Per paradigm, single deflections were carried out at an inter-stimulus interval of 20 seconds, for 20 repetitions.

### Analysis of *in Vivo* Multi-Electrode Silicon Probe Data

Extracellular recordings were analyzed using custom made MatLab script (R2010a, MathWorks). For each experiment the channel(s) on a shank were positioned within an intrinsically located barrel of vS1. The depth of the recording channels were assessed by leaving the top channel of each shank outside the cortical plate as well as creating vertical current source density maps (CSD) (Reyes-Puerta et al. 2015). CSD maps were created from the LFP averages, gathered while performing the whisker stimulation paradigms. The shanks position allowed for identification of L1 while the early onset and shape of the CSD response after whisker deflection allowed for identification of the recipient thalamo-cortical L4, as well as then L2/3 and L5 (Mitzdorf 1985, van der Bourg et al. 2017). For MUA registration, the signal of each electrophysiological recording was bandpass filtered at 800-5000 Hz (Reyes-Puerta et al. 2015, van der Bourg et al. 2017). A threshold of either 6.5 or 3.5 standard deviations from baseline was applied and the time stamp of detected MUA was registered. The MUA were separated per stimulation paradigm and peri-stimulus time histograms (PSTHs) were created. PSTHs were made from the summation of activity from all individual trials recorded within group, and MUA ms^-1^ was normalized to number of trials. The PSTHs were plotted as CTRL vs. KO and analysis were done within three temporal domains; between 5-35 ms (*early*), 35-100 ms (*late*) and 100-300 ms (slow) after stimulus onset, presented as dashed line at 0 ms.

### Analysis of *in Vivo* Whole-Cell Patch-Clamp Data

To remove any electrical noise, a 50 Hz Notch filter was applied to all data using Clampfit (v10.7.0.3, Molecular Devices LLC.). The whisker evoked responses belonging to each stimulation paradigm and mice group was extracted using custom made Python 3.9.0 script. The responses were separated per recorded voltage-clamp mode and a baseline (average of 10 ms period before stimulus delivery) was subtracted from the signal. Average recorded activity for all trials within specific stimulation paradigm were plotted as CTRL vs. KO. For visualization purposes only, a rolling average of 15 was applied.

### Statistical analysis

Data represented as averages ± S.E.M. unless otherwise stated. Statistical comparisons carried out by using a paired Wilcoxon-signed rank test for paired data and two-tailed Mann-Whitney U test for non-paired data. Significance threshold was set to p<0.05; in the figures, different degrees of evidence against the null hypothesis are indicated by asterisks (p<0.05: *;p<0.01: **;p<0.001: ***).

## Acknowledgements

We thank O. Hanley for the production of the pseudotyped rabies virus and the HTB plasmid used in this study, G. Kanatouris for optimizing the NPY antibody staining, J.W. Yang for the teaching of *in vivo* silicon probe recordings, H. Kasper for technical assistance. Slidescanner imaging and data analysis was performed with equipment maintained by the Center for Microscopy and Image Analysis (ZMB), University of Zürich. This work was supported by grants from the European Research Council (ERC, 679175, T.K) and the Swiss National Science Foundation (SNSF, 31003A_170037,T.K).

## Author contributions statement

R.V. performed; viral injections, morphological reconstructions, anatomical mapping, *in vitro* electrophysiology experiments, *in vivo* multi electrode experiments, the analysis of the *in vitro* electrophysiology data, the functional *in vivo* patch-clamp data as well as the multi electrode data, contributed to the *in vivo* patch-clamp experiments and wrote the manuscript; L.G. bred and maintained the mouse lines, validated the Ndnf-IRES2-dgCre-D, contributed to the validation of the NDNFCre × NR1 ^*floxed*^ knockout, fate-mapped the NDNF line and performed confocal imaging; A.D. performed viral injections, anatomical mapping, immunohistochemistry, confocal imaging, slide scanner imaging and quantification of imaged data; L.B. performed the *in vivo* patch-clamp experiments; V.S. performed the analysis of the functional *in vivo* and *in vitro* electrophysiology data; T.K. conceptualized the study and wrote the manuscript.

## Additional information

### Data availability

Data are available upon request.

### Accession codes

Codes written for this manuscript are available upon request.

---

**Supplemental Figure 1:**
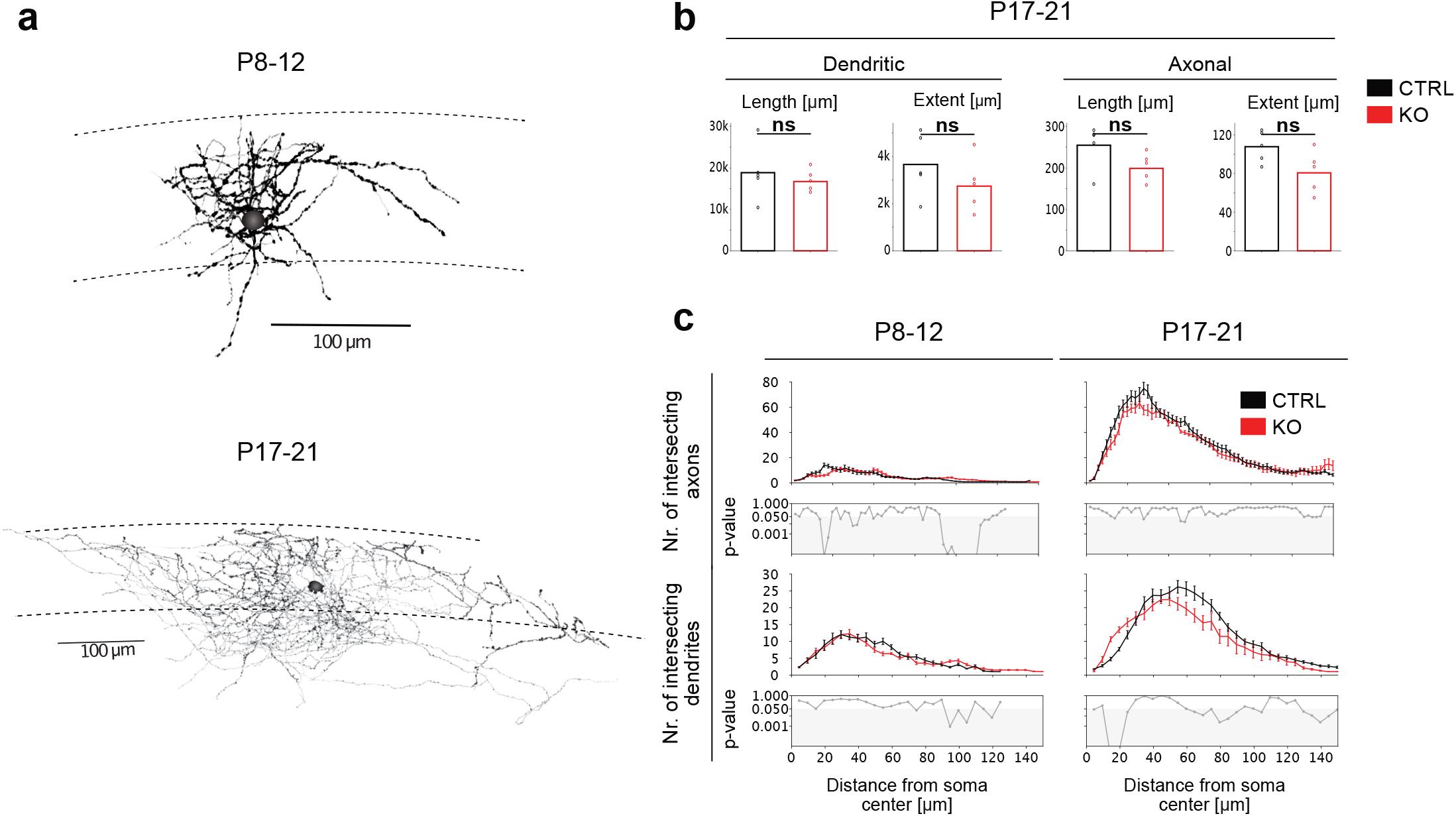
Maintained physiological morphology within NDNF^+^vS1 neurons both at P8-12 and P17-21, with a failure of integrating evoked EPSPs, when selectively removing the NMDA receptor. **(a)** Two example NDNF^+^cells filled with biocytin in vitro before and after P14 and subsequently morphologically reconstructed. **(b)** No significant difference in the total length (length of all combined parts of neurite) or extent (distance from cells soma to end of furthest away neurite) between dendrites or axons of morphologically reconstructed vS1 L1 NDNF^+^ CTRL or KO neurons (CTRL Dendrites: 5 cells, KO Dendrites: 5 cells; CTRL Axons: 5 cells, KO Axons: 5 cells). **(c)** Plotted overall average of sholl analysis data from all neurites of morphologically reconstructed neurons. Number of intersecting neurites plotted per distance step of 5 *μ*m from the cells soma, with statistical comparison performed for all cell averages per step distance (CTRL P8-12: 5 cells, KO P8-12: 5 cells; CTRL P17-21: 5 cells, KO P17-21: 4 cells). Statistics: two-tailed Mann-Whitney U-test (*p<0.05, **p<0.01, ***p<0.001).

